# MiR-219 deficiency in Alzheimer’s disease contributes to neurodegeneration and memory dysfunction through post-transcriptional regulation of tau-kinase network

**DOI:** 10.1101/607176

**Authors:** Mercedes Arnes, Yoon A. Kim, Jerome Lannes, Maria E. Alaniz, Joshua D. Cho, Brian D. McCabe, Ismael Santa-Maria

## Abstract

Intracellular accumulation of hyperphosphorylated misfolded tau proteins is one of the main neuropathological hallmarks in Alzheimer’s disease (AD) and related tauopathies. Hence, knowledge and understanding of disease mechanisms altering tau proteostasis and inducing cytotoxicity is critical. MicroRNAs (miRNAs) are capable of binding to and silencing many target transcripts, providing an additional level of regulation that complements canonical transcriptional pathways. Therefore, observed abnormalities in their expression patterns in neurodegeneration suggest alterations of microRNA-target networks as drivers of cellular dysfunction in the disease. Strikingly, here we have found in autopsy brain tissue that miRNA miR-219 expression levels are decreased in a brain region early affected in AD patients, the entorhinal cortex. Our bioinformatics analysis indicates miR-219 is predicted to target Calcium/calmodulin-dependent protein kinase 2 gamma subunit (CAMK2γ), Tau tubulin kinase 1 (TTBK1) and Glycogen synthase kinase 3 beta (GSK3β), which are all implicated in the generation of abnormal hyperphosphorylated tau. We reveal human proteomic data supporting dysregulation in the levels of predicted miR-219 targets in the entorhinal cortex. In mammalian cellular models, we found that downregulation of miR-219 de-repress synthesis of three tau kinases, CAMK2γ, TTBK1 and GSK3β on the post-transcriptional level resulting in tau phosphorylation and cell toxicity. Finally, we show that deficiency of miR-219 *in vivo* promotes age dependent neurodegeneration in the adult brain, with enhanced alterations in tau proteostasis, presynaptic terminals and memory impairment. Taken together, our data implicate miRNA dysregulation central to AD etiopathogenesis and suggest potential targets for the treatment of AD and related tauopathies.

## Introduction

Alzheimer’s disease is a neurodegenerative disorder involving complex cellular and molecular networks that over time alter the homeostasis of the brain in specific areas driving massive synaptic loss and atrophy^1,2^. MicroRNAs (miRNAs) are powerful key regulators of gene expression contributing to overall health, molecular homeostasis and neuroprotection of the brain^3^. However, in AD the expression of miRNAs is dysregulated suggesting a functional involvement in the pathogenic process, even though it remains largely unknown how these aberrations are induced and whether they play a role in disease progression. Only a small number of miRNAs have been found consistently and robustly altered in AD brains^4,5^. One of the few miRNAs that are consistently downregulated in the disease is miR-219. MiR-219 is a neuroprotective and highly conserved ancient microRNA enriched in the mammalian nervous system^6,7^ that play essential roles in a myriad of molecular and cellular processes including neural precursor differentiation, myelination, metabolic responses, inflammatory pain, NMDA receptor regulated neurotransmission, circadian regulation and synaptic plasticity^8–15^. Moreover, we have previously described miR-219 role in the posttranscriptional regulation of tau synthesis^5^. All these miR-219 roles suggest dysregulation of this miRNA could play a central role in the pathogenesis of AD.

In this study, for the first time we have determined miR-219 expression levels are decreased in the entorhinal cortex of AD patients, a brain region affected in early stages of the disease. Conversely, the protein levels of the three predicted miR-219 targets, CAMK2γ, TTBK1 and GSK3β are found upregulated in this brain region in AD. Our results show miR-219 loss alters the synthesis of CAMK2γ, TTBK1, and GSK3β, leading to alterations in tau expression and tau hyperphosphorylation supporting a unifying mechanism that drives tau toxicity in the disease through alterations in specific molecular networks. Further, we have characterized the functional impact miR-219 deficiency has on AD-tau pathology *in vivo*. MiR-219 deficiency *in vivo* aggravates tau pathology, neurodegeneration and impairs learning and memory. Taken together, our data implicate miR-219 dysregulation central to AD pathogenesis and suggest miR-219 as potential target for the treatment or prevention of AD and related tauopathies.

## Results and Discussion

Previous studies have shown in mice adult forebrain neurons, genetic ablation of Dicer, a key enzyme in the biogenesis of miRNAs, results in abnormal tau hyperphosphorylation, tau accumulation and neurodegeneration^16^. Thus, providing a proof of concept that miRNA dysregulation causes abnormalities in tau proteostasis. Given a single miRNA is capable of binding to and silencing several targets this opens up the possibility that dysregulation of particular miRNAs in AD induce changes in expression of tau and specific kinases in synchrony (Figure 2A, Supplementary Figure 1). We have previously shown miR-219 regulates tau synthesis on the post-transcriptional level supporting alterations in the levels of this particular miRNA are involved in the progression of AD^5^. Hence, we investigated whether alterations in miR-219 are observed in the brain region of the human brain that is primarily affected in patients with AD. We found miR-219 levels are decreased significantly in the entorhinal cortex of the AD brains when compared to non-demented controls (Figure 1A, Supplementary Table 1). Importantly, we could determine miR-219 downregulation is significant in cases with sparse-moderate pathology supporting a possible role for miR-219 in the onset of the disease (Figure 1B).

**Figure 1.**
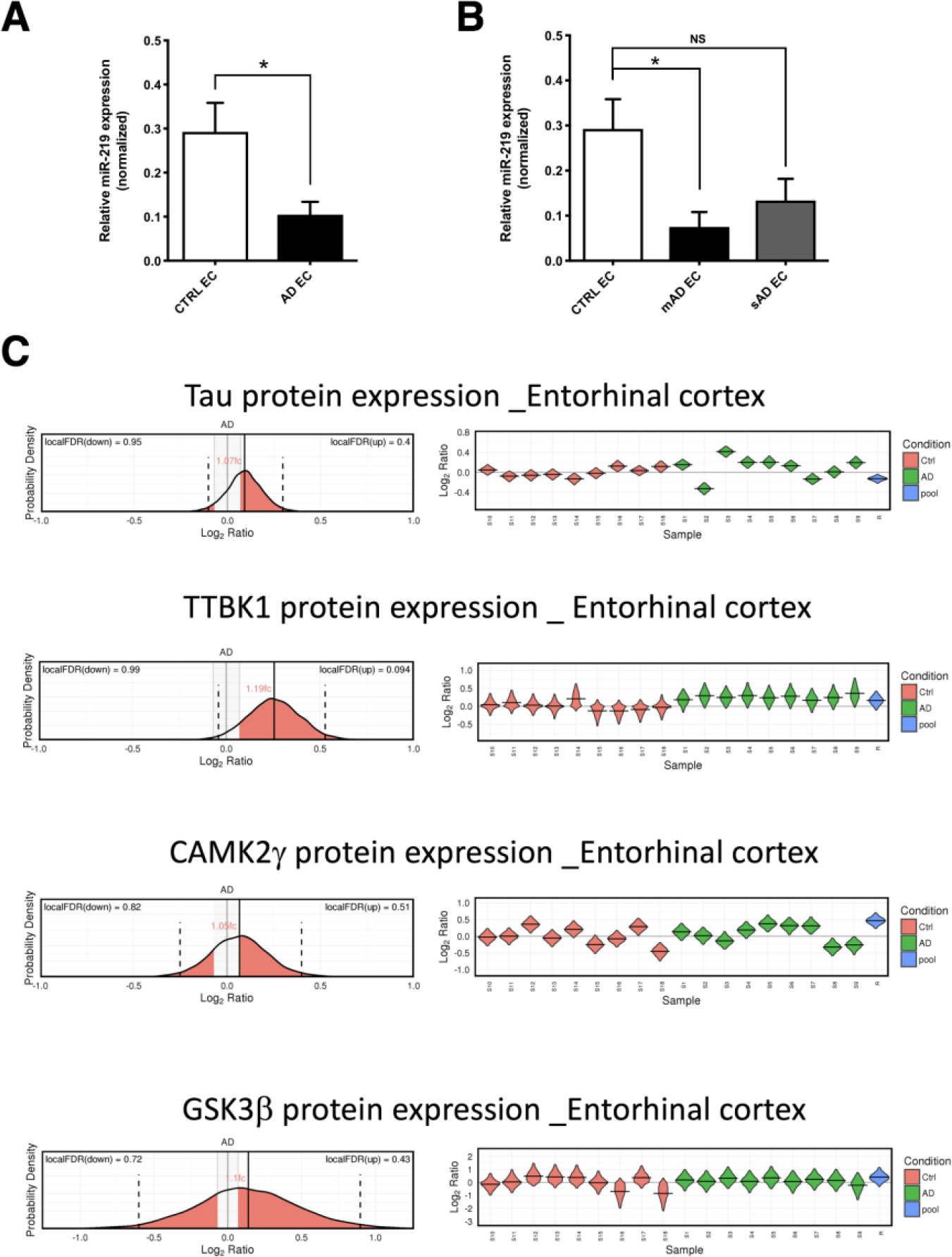
Alterations in the levels of miR-219 and miR-219 targets in the human entorhinal cortex with Alzheimer disease. **(A)** qPCR confirmed decreased levels of miR-219 in AD entorhinal cortex (EC) (n=8) postmortem brain tissue samples compared with those of controls (CTRL EC) (n=4). **(B)** miR-219 expression levels determined on control, moderate (mAD) and severe (sAD) AD cases by qPCR. U6 snRNA levels were used for normalization. *P ≤ 0.05 by 2-tailed Student’s t test (Supplementary Table 2). **(C)** Probability density plots for miR-219 targets Tau, TTBK1, CAMK2γ, GSK3β and CDK5 (a kinase not targeted by miR-219) from entorhinal cortex AD proteomic dataset is shown. Algorithm used by Xu et al ^27^ (http://www.dementia-proteomes-project.manchester.ac.uk/Proteome/Search) calculates a local FDR (1-p (protein differs from control by at least 5%)) both for upregulation and downregulation. Mean Fold change (fc) is indicated in the AD density plot for each represented protein. Variability across all entorhinal cortex samples (Alzheimer disease and controls) is shown on the right of every Probability density plot shown for each protein represented.

**Figure 2.**
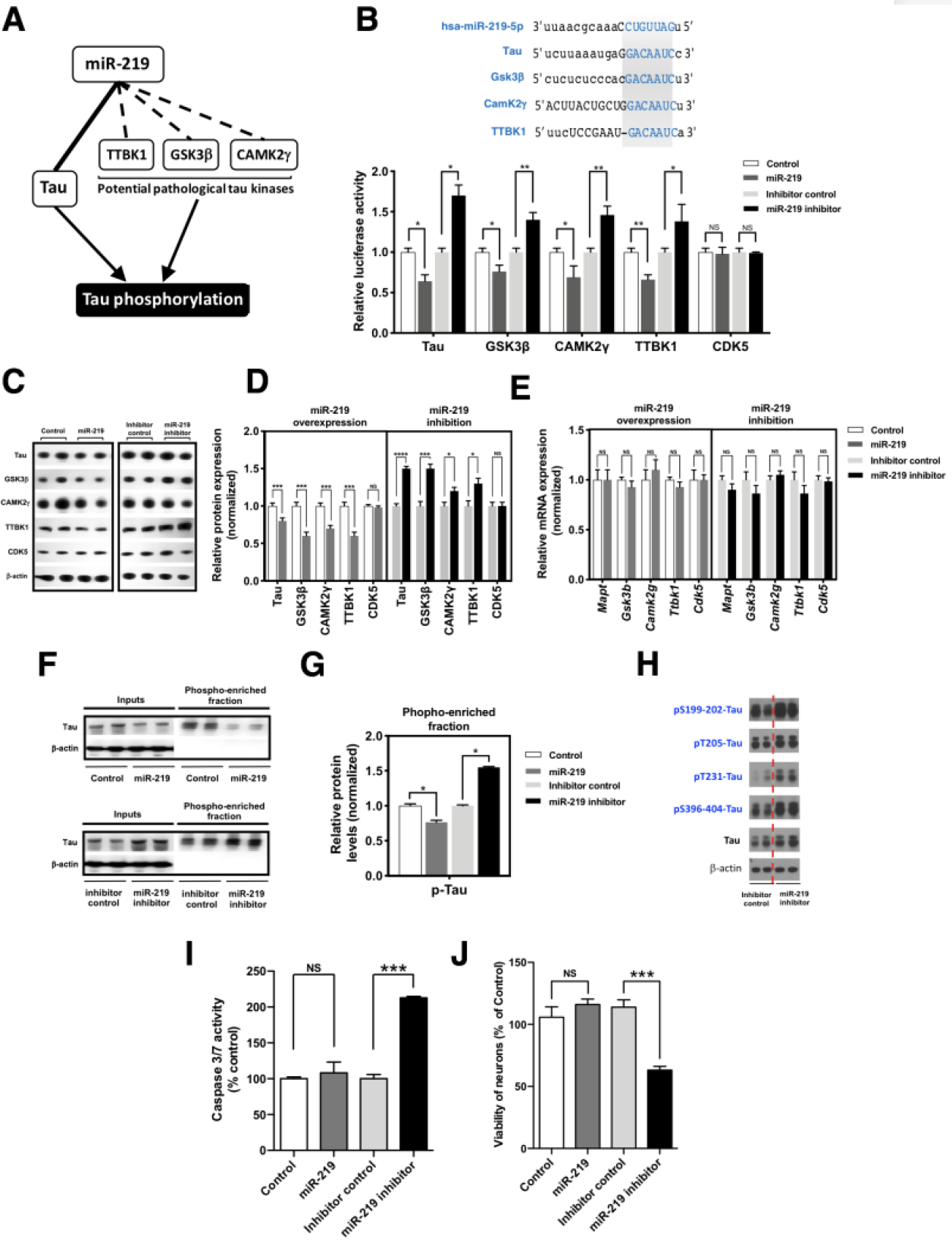
MiR-219 attenuates tau hyperphosphorylation and directly regulates expression of tau kinases GSK3β, CAMK2γ and TTBK1. **(A)** Schematic illustrating synchronous regulation of tau and tau kinases by miR-219. (**B**) Both the tau & tau kinases (TTBK1, GSK3β, and CAMK2γ) 3’ UTR have highly conserved miRNA recognition elements for miR-219. Relative luciferase activity of Tau, GSK3β, CAMK2γ, TTBK1, CDK5 (negative control) reporter constructs co-transduced with miR-219 or miR-219 inhibitor-expressing vector and their respective controls in human SH-SY5Y cells (n=3 with 6 replicates each, mean ± SEM, \**P*<0.05, \*\**P*<0.01, Student’s *t*-test. (**C, D**) Immunoblots with indicated antibodies and quantification after normalization to β-actin levels, n = 3, mean ± SEM, \**p*< 0.05, \*\*\**P*< 0.001, \*\*\*\**P*< 0.0001, Student’s *t* test. (**E**) The mRNA levels for studied miR-219 targets are unchanged as assessed by QPCR. (**F-G**) Immunoblots with indicated antibodies and quantification of phospho-enriched fractions from primary neuronal cultures transduced with lentiviral miR-219 or miR-219 inhibitor and their respective controls (n=3), mean ± SEM, \**P*<0.05, Student’s *t*-test. (**H**) Immunoblots with indicated antibodies against phospho-tau shows specific sites are increased after lentiviral expression of miR-219 inhibitor and the inhibitor control in primary neuronal cultures. (**I**) Caspase activity was assessed using the Caspase-3/7 Glo kit on primary neuronal cultures transduced with lentiviral miR-219 or miR-219 inhibitor and their respective controls. Mean ± SEM, n=6, \*\*\**P*<0.001, Student’s t-test. (**J**) Cell viability was measured using the Cell Titer Glo kit on primary neuronal cultures transduced with lentiviral miR-219 or miR-219 inhibitor and their respective controls. Mean ± SEM, n=6, \*\*\**P*<0.001, Student’s t-test.

Pinning down the relevant tau kinase is challenging as tau is phosphorylated on over 20 sites and no single kinase can phosphorylate tau on all these residues^17^, suggesting multiple kinases work in concert to generate hyperphosphorylated tau. Using computational algorithms for microRNA target prediction^18^ on all known tau kinases (i.e. Cdk5, GSK3β, MARK, PKA, AMPK)^17,19,20^ revealed only three kinases are predicted targets for miR-219 and highly conserved in both humans and rodents (mouse and rat), Tau tubulin kinase 1 (TTBK1), Calcium/calmodulin-dependent protein kinase 2 gamma subunit (CAMK2γ) and Glycogen synthase kinase 3 beta (GSK3β) (Supplementary Figure 1), which are all implicated in the generation and/or accumulation of abnormal hyperphosphorylated tau^21–25^. Importantly, miR-219 is not predicted to bind to the 3’UTR and repress expression of any known tau phosphatases^26^.

This finding prompted us to explore the AD brain proteome^27^. We found alterations in the protein expression of the three miR-219 predicted targets, TTBK1, CAMK2γ and GSK3β in the human entorhinal cortex of brains with AD (Figure 1C). However, we didn’t observe mRNA levels alterations for any of these kinases in the human brain tissue we assessed (Supplementary Figure 2, Supplementary Table 2), supporting alterations on the posttranscriptional regulation of the three kinases.

We next confirmed that miR-219 silences the expression of the three candidate kinases, TTBK1, CAMK2γ and GSK3β in human cells and asked whether this occurs through a direct interaction with the 3′-UTR. The full-length 3’ UTR of the three candidate kinases, TTBK1, CAMK2γ and GSK3β was inserted into a dual-luciferase reporter construct downstream of the Renilla luciferase. We used as a negative control the CDK5-3’UTR that does not contain any predicted miR-219 binding element. We transiently co-transfected these constructs along with lentiviral transduction of miR-219 or scrambled control into cell cultures. We found that miR-219 significantly reduce luciferase activity of all the three candidate kinases but does not reduce CDK5-3’UTR activity when compared with that in the control, while transduction with a specific miR-219 inhibitor abrogated silencing when compared to an inhibitor control (Figure 2B). These findings demonstrate that silencing of TTBK1, CAMK2γ and GSK3β expression by miR-219 occurs through a direct interaction with the predicted and highly conserved recognition elements (Supplementary Figure 1) in the candidate’s kinases 3’ UTRs. Further, we confirmed miR-219 post-transcriptionally regulates candidate kinases in human neuronal cell cultures. Lentiviral transduction of miR-219 resulted in a statistically significant reduction in the protein levels of all the candidate kinases when compared to scrambled control (Figure 2C, 2D). Interestingly, QPCR analysis did not show a difference in candidate kinases mRNA levels (Figure 2E), suggesting that, in this context, the silencing mechanism of miR-219 is through translational repression. In addition, depletion of miR-219 levels with a lentiviral miR-219 inhibitor resulted in a significant increase in candidate kinases protein levels compared with those of the negative control kinase, CDK5 (Figure 2C, 2D). Similarly, expression of the miR-219 inhibitor did not resulted in a significant increase in candidate kinases mRNA levels (Figure 2E). These findings support alterations in miR-219 levels induces changes in the synthesis of miR-219 targeted kinases.

To demonstrate alterations in miR-219 levels modulate tau hyperphosphorylation and neurotoxicity first we have confirmed transduction of lentiviral miR-219 inhibitor in primary neuronal cultures induces a significant increase in phosphorylated tau levels as observed by phospho-enrichment analysis (Figure 2F-G). MiR-219 inhibition increases levels of certain phospho-tau epitopes phosphorylated by TTBK1, CAMK2γ and GSK3β (Tau pSer199-202, Tau pThr205, Tau pThr231 and Tau pSer396-404; http://cnr.iop.kcl.ac.uk/hangerlab/tautable) as observed by western blot using phospho-tau specific antibodies (Figure 2H). Having observed miR-219 regulates the three tau kinases on the post-transcriptional level we then explored whether miR-219 inhibition could promote cytotoxicity. Remarkably, miR-219 downregulation induced cleavage of Caspase 3/7 and reduced neuronal survival significantly (Figure 2I, 2J). Our observations collectively support miR-219 dysregulation alters the synthesis of GSK3β, CAMK2γ, TTBK1 leading to tau hyperphosphorylation and cell toxicity.

To investigate the effect miR-219 deficiency has *in vivo* we took advantage of a previously generated miR-219 mutant knockout *Drosophila* line obtained by targeted homologous recombination^28^ (Supplementary Methods). In order to determine whether loss of miR-219 plays a role in degeneration of the adult brain, we first tested whether miR-219 deficiency would promote toxicity in an age-dependent manner in miR-219 mutant Drosophilae. Loss of miR-219 in the *Drosophila* brain results in a progressive neurodegenerative phenotype characterized by histological abnormalities, including neuropil vacuolization and degeneration of brain cells in the cortex^29–31^. Processes of brain cells are located in the inner neuropil region, which is eosinophilic and stains pink or light purple following H&E processing. The cell bodies of brain cells occupy the outer cortical layer of the brain, which is basophilic and can be distinguished after H&E staining by a dark purple color (Figure 3A-C and Supplementary Methods). MiR-219 loss enhanced toxicity in the brain as evidenced by an increase in neuropil vacuolization (Figure 3A-F). The brains of control flies were well preserved at 7, 20 and 45 days of age, but we observed neuropil vacuolization in miR-219 knockouts flies by 20 and 45 days of age (Figure 3A-F). Quantification of the total number of vacuoles present in the neuropil of 7, 20 and 45-day-old transgenic animals and controls revealed that brain vacuolization in miR-219 knockout flies was significantly increased by 2-fold in 20 and 40 days old miR-219 mutants, compared with that of control animals (Figure 3D-F). The increased presence of the active forms of caspases has been observed in Alzheimer’s brain relative to controls^32–34^. To further investigate the mechanism of brain neurodegeneration, we performed immunofluorescence using antibodies against activated caspase 3, an effector of apoptosis. We found activated caspase staining within the cytoplasm of brain cells in the cortex of 20 and 45 days old miR-219 mutants flies which was enhanced significantly in an age dependent manner when comparing to control flies (Figure 3G, 1H), suggesting an apoptotic mechanism of cell death. To investigate the extent to which cellular damage in the cortex brain cells accompanied degeneration observed in the neuropil of miR-219 mutants we assayed control and miR-219 mutant flies for the presence of TUNEL-positive nuclei (Figure 3I, 3J) to mark fragmented DNA within nuclei of damaged brain cells^29,30^. We performed fluorescence TUNEL labeling on 20, and 30-day-old control and miR-219 flies and observed a significant age-dependent increase in TUNEL-positive cells (Figure 3I, 3J). Curiously, despite all the degenerative changes observed in the miR-219 mutant flies, there was not a significant reduction in the lifespan when compared to control flies (Supplementary Figure 3). We have previously shown miR-219 regulates tau expression at the post-transcriptional level through direct interaction with the tau mRNA^5^. The accumulation of sarkosyl-insoluble species of tau is a defining characteristic of tau isolated from brains of patients with Alzheimer’s disease and related tauopathies^35,36^. Next, we asked whether loss of miR-219 promotes alterations in tau proteostasis *in vivo*. First, we were able to confirm in the fly adult brain miR-219 dysregulation alters the levels of tau protein but not tau mRNA suggesting effects on mRNA translation (Figure 4A-C). Further, we performed western blot analysis on isolated tau sarkosyl insoluble and soluble fractions from fly head homogenates of 20-day-old miR-219 mutants and aged-matched control flies. As expected, we did detect the presence of tau in the sarkosyl-soluble homogenates of 20-day-old miR-219 mutant and control flies (Figure 4D, lower panel). Strikingly, we did observe an increase in the amount of sarkosyl-insoluble tau recovered from 20 days old miR-219 mutant flies (Figure 4D, upper panel). Moreover, when miR-219 mutant flies were crossed with a line with the endogenous *Drosophila* tau tagged with GFP we were able to visualize accumulation of endogenous tau in the fly brain in an age dependent manner (Figure 4E, Supplementary Video 1, Supplementary Video 2).

**Figure 3.**
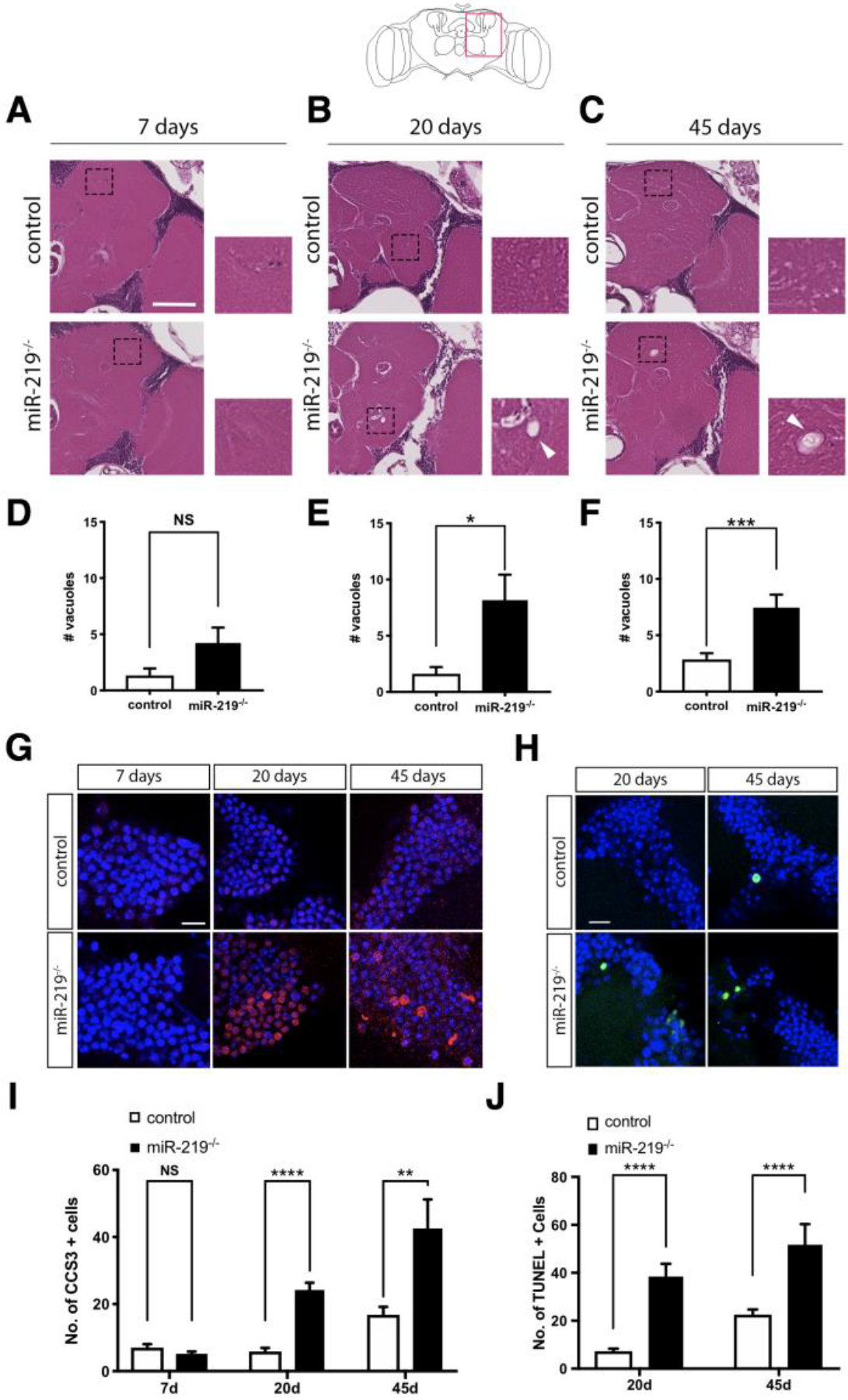
MiR-219 loss promotes age-dependent neurodegeneration in the adult brain of Drosophila. Representative frontal sections from adult *Drosophila* brains at 7 days (**A**) 20 days (**B**) and 45 days (**C**) of control (*w*^1118^) and miR-219^−/−^ flies stained with hematoxylin and eosin staining. High magnification images of the corresponding dotted areas show vacuole presence (arrowheads). (**D**) Neurodegeneration was not observed in control flies and miR-219^−/−^ flies at 7 days. Quantification of vacuole numbers in 20 (**E**) and 45 (**F**) days old miR-219^−/−^ *Drosophila* brains shows a significant increase in vacuole presence when compared to control flies. **(G)**Representative confocal images for activated caspase 3 (red) and DAPI (blue) immunostainings in adult *Drosophila* brains of 7, 20- and 45-days old *Drosophila* brains. Scale bar is 10μm. (**H**) Quantification of the number of activated Caspase-3 positive cells through entire brains of control and miR-219^−/−^ flies. MiR-219^−/−^ flies show increased number of activated caspase-3 positive cells in 20 days and 45 days brains. (**I**) Representative images of adult fly brains immunofluorescence TUNEL assay in control and miR-219^−/−^ flies at 20 and 45 days. Scale bar is 10 μm. (**J**) TUNEL-positive cells (green) quantification through the entire brain showing a significant increase in the number of TUNEL-positive cells in miR-219^−/−^ flies of 20 days and 45 days. Mean ± SEM, \**P*<0,05; \*\**P*<0,01 and \*\*\**P*<0,001 by Mann-Whitney U test.

**Figure 4.**
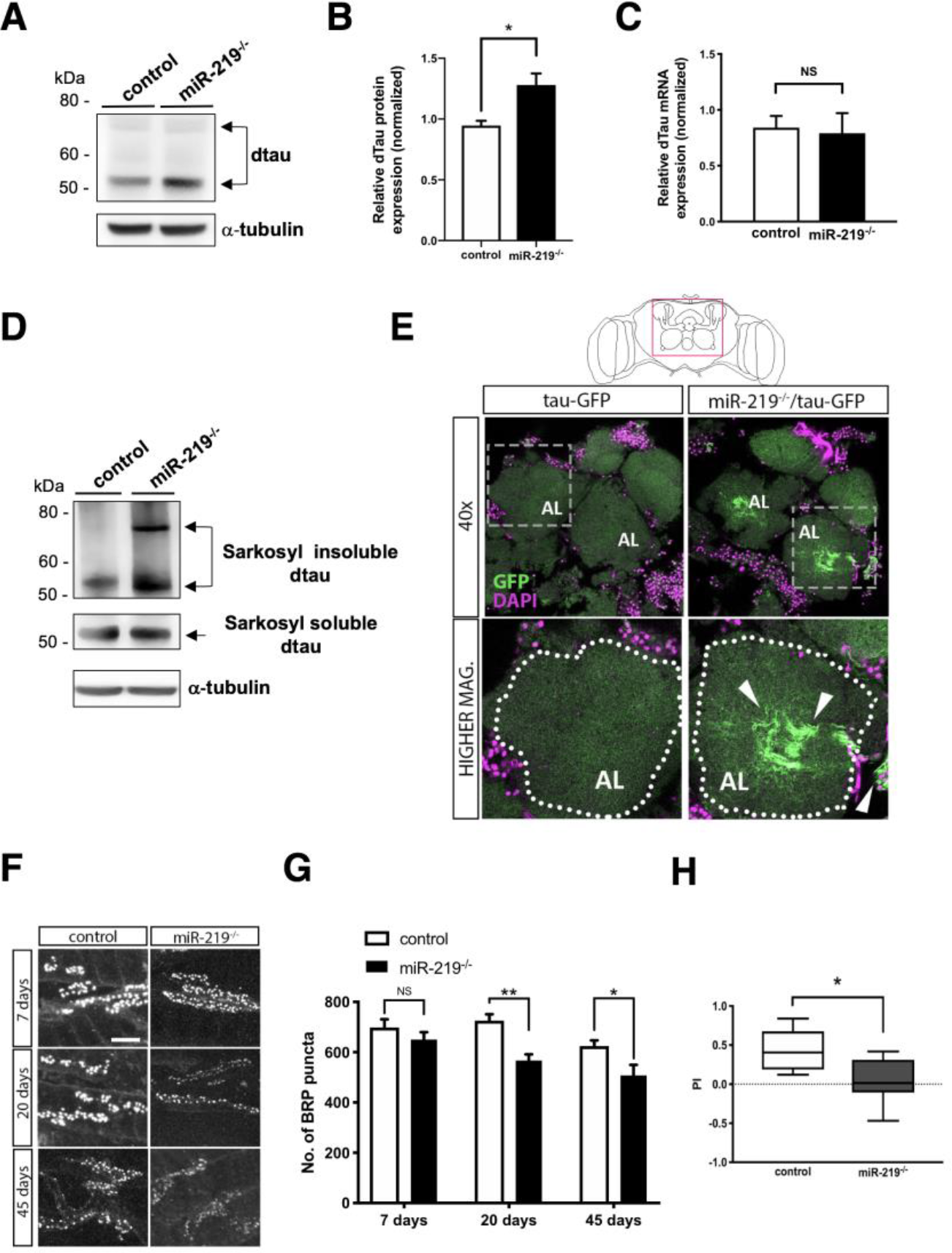
MiR-219 loss promotes accumulation of insoluble tau, alterations in presynaptic terminals and memory deficits. **(A-B)** Immunoblot with tau antibody and quantification after normalization to α-tubulin levels of Drosophila head lysates from control and miR-219^−/−^ flies. Mean ± SEM, \**P*<0.05, Student’s *t*-test. **(C)** QPCR showing unchanged levels of mRNA dtau levels in miR-219^−/−^. **(D)** Immunoblot of sarkosyl extracted brain homogenates showing increased dtau levels both in the sarkosyl insoluble and soluble protein fractions (n=3). **(E)** Brain confocal images of control and miR-219^−/−^ flies expressing GFP-tagged tau showing tau accumulation throughout the brain. Mutant flies lacking miR-219 showed enhanced tau accumulation in different areas of the brain, both in the somas and in the neuropil region (arrowheads) (n=3). (AL: antennal lobe). **(F)** High-magnification confocal images of adult NMJs from control and miR-219^−/−^ flies of 7, 20 and 45 days showing BRP-positive puncta. Scale bar is 5 mm. Full images are shown in Supplementary Figure 4. **(G)** Synapse number quantification of the corresponding genotypes showing number of BRP-positive puncta in NMJs of the adult ventral longitudinal muscle. Data show that the number of synapses is decreased in miR-219^−/−^ flies at 20 days and 45 days comparing to age-matched controls (n=6-8/group). **(H)** Performance Index of aversive associative memory assay reporting a significant decrease in miR-219 mutant flies (n = 400/group). Mean ± SEM, \**P*<0.05, \*\**P*<0.01 Mann-Whitney U test.

Pathological tau was previously detected at synaptic compartments of neurons in Alzheimer’s disease. Several studies support tau triggers pathophysiology in the brain by altering properties of synaptic and neuronal function at the early stages of the disease, before neuronal death becomes evident^37–46^. We then investigated whether miR-219 loss promotes alterations in synaptic terminals in the *Drosophila* neuromuscular junctions (NMJs), a large, accessible synapse which can be analyzed during all stages of its development and in adult flies. Given that *Drosophila* synapses develop and function similarly to the synapses of higher mammals using conserved genetic and molecular mechanisms^47^ and that is been described pathological tau causes NMJ presynaptic damage in *Drosophila*^46^ we evaluated the active zones at synaptic terminals of the *Drosophila* NMJ by confocal microscopy using an antibody that recognizes the NMJ presynaptic marker Bruchpilot (BRP)^48,49^. We found that the active zone number is decreased in the miR-219 mutant adult flies in an age-dependent manner when compared to control flies. A significant reduction of the number of BRP puncta was observed in miR-219 mutant flies at 20 and 45 days but not as early as 7 days (Figure 4F, 4G; Supplementary Figure 4), correlating with overall neurodegeneration observed in the brains of these miR-219 mutant flies (Figure 3). Strikingly, synaptic changes were not observed in larvae stages (Supplementary Figure 5) supporting the role miR-219 plays in the adult brain and neurodegeneration.

It has been proposed elevated hyperphosphorylated tau in the “pre-tangle” (non-neurofibrillary tangle) state observed in tauopathies may underlie pre-neurodegeneration cognitive symptoms including memory loss^50,51^. Therefore, we decided to investigate whether learning and memory was affected in the miR-219 mutant flies which show tau proteins accumulating in their brains. We found that miR-219 deficiency precipitated a significant decrement in associative learning and memory retrieval or stability as early as 7 days in miR-219 mutant flies compared to controls (Figure 4H). Indicating early cognitive symptoms are found, as in humans, in miR-219 mutant flies.

In conclusion, these results support that miR-219 dysregulation promotes age-dependent neurodegeneration and memory alterations by modulating tau pathology through direct regulation of synthesis of not only tau but also tau kinases, TTBK1, CAMK2γ and GSK3β. We based this conclusion on the obtained results showing miR-219 specifically binds the 3′-UTR of tau and tau kinases *in vitro* influencing tau and tau kinases expression at the post-transcriptional level, mainly through translational repression. Given the extraordinary and unique conservation of miR-219 recognition elements in the tau and tau kinases 3′-UTRs extended validation *in vivo* will be of great value. Further studies will advance our understanding of the pathogenesis of neurofibrillary degeneration and may guide us toward the development of novel therapeutic strategies.

## Methods

Detailed Supplementary Methods are available online

### Statistical analysis

Statistical significance was calculated using a Mantel-Cox test for lifespan analysis, and unpaired Mann-Whitney test for the rest of the experiments involving Drosophila, with significant differences between compared groups noted by \**P*≤0.05, **P≤0.01, and ***P≤0.001.

The statistical significance was determined using Prism (GraphPad Software). For all figures in which error bars are shown, data are means ± SEM. Statistical outliers and specimens with measurement errors were

### Study approval

The studies using de-identified post-mortem autopsy tissue were reviewed and approved by the Columbia University institutional review board (IRB) (New York, NY). Drosophila studies are not subject to IRB oversight.

## Acknowledgments

This work was supported by the National Institute of Neurological Disorders and Stroke (R01NS095922) and the National Institute of Aging (P50AG0008702). Additional support was provided by the Alzheimer’s Association (NIRG-15-3644-58). We want to thank the Columbia University College of Physicians and Surgeons office for research for critically reading the manuscript, and Michael L. Shelanski, Lloyd A. Greene and Carol M. Troy for comments and discussions. We also want to thank Daisuke Hattori, Richard Axel and James Hodge for their help and advice on *Drosophila* behavior experiments. Finally, we thank Etty Cortes, Jean Paul Vonsattel and Andrew F. Teich for neuropathology support.

