## Supplementary materials for "MiR-219 deficiency in Alzheimer’s disease contributes to neurodegeneration and memory dysfunction through post-transcriptional regulation of tau-kinase network"

### This file includes

Supplementary Methods

Supplementary Figures 1 to 5

Supplementary Videos 1 and 2

Supplementary Tables 1 to 3

Supplementary References

**Methods**

*Patient samples.* De-identified human autopsy brain tissue (Supplementary Table 1, 2) was obtained from the New York Brain Bank at Columbia University.

*RNA isolation.* Fresh-frozen brain tissue was pulverized in liquid nitrogen, lysed in QIAzol and homogenized using a QIAshredder column. Total RNA enriched in microRNAs was extracted from human brain tissue or cell cultures using the miRNeasy Kit (Qiagen).

*Quantitative Real-Time PCR (QPCR). c*DNA synthesis was performed using the First Strand cDNA Synthesis Kit (Origene). QPCR was performed on a Mastercycler ep realplex (Eppendorf) using TaqMan Gene Expression Master Mix and primers and TaqMan probes specific for total tau, tau kinases and GAPDH mRNAs (Life Technologies). The following settings were used: 95 °C for 10 min followed by 40 cycles of 95 °C for 15 s and 60 °C for 1 min. The GAPDH mRNA levels were used for normalization. TaqMan MicroRNA assays were used to measure miR-219-5p levels (Life Technologies). 100 ng of total RNA was reverse-transcribed using specific stem-loop reverse transcription primers (Life Technologies) and miR-219-5p levels were measured on a Mastercycler ep realplex (Eppendorf). The levels of U6 snRNA were used as endogenous controls for normalization using the comparative CT method.

*Target prediction.* Currently, integration of various computational methods is a common approach to improve prediction accuracy and to create an optimal framework for deciphering biological functions of miRNAs[1]. We used TargetScan[2] and miRBase[3] to identify predicted kinases 3' UTR targets for miR-219 (Supplementary Figure 1). TargetScan is a well-established algorithm of seed and sequence complementarity with conservation of binding sites across multiple species that has been shown to result in the most accurate predictions upon target validation[4,5].

*Lentiviral vectors.* The human mir-219 precursor, miRNA control, miR-219 inhibitor and inhibitor control were obtained from Applied Biological Materials Inc. Lentiviral production was performed as described by others[6]. We used the human HEK 293T cell line for optimal lentivirus production. HEK 293T cells (ATCC) were grown either in Dulbecco's modified Eagle's medium (DMEM) or DMEM/F12 medium (Cellgro) supplemented with 10% fetal bovine serum, 2 mM glutamine, 100 units/ml penicillin, and 100 μg/ml streptomycin in a humidified atmosphere of 5% CO2 and 95% air at 37 °C. Lentiviral stock titration was carried out using the Global UltraRapid Lentiviral Titer Kit (System Biosciences).

*Cell culture.* Human SH-SY5Y cells (ATCC) were grown either DMEM/F12 medium (Cellgro) supplemented as mentioned above for HEK 293T cells.

*Luciferase assays*. For 3′ untranslated (UTR) reporter assays, CalFectin™ Mammalian Cell Transfection Reagent (SignaGen Laboratories) was used to co-transfect human SH-SY5Y cells (ATCC) with 3′ UTR target luciferase reporter vectors for human TTBK1, CAMK2γ, GSK3β, CDK5 (Genecopoeia), and mir-219 precursor, miRNA control, miR-219 inhibitor and inhibitor control in 96-well plates. Cells were lysed 48 h post-transfection, and luciferase activities were measured using the Genecopoeia Luc-Pair miR Luciferase Assay Kit as instructed by the manufacturer using an Infinite F200 plate reader (TECAN).

*Primary neuronal cultures.* Primary cortical neuronal cultures were prepared from embryonic day 18 rat embryos following previously described methods[7]. Cortical neurons were plated on poly-d-lysine-coated culture plates in Neurobasal medium containing B27 supplement and 0.5 mM glutamine (Invitrogen).

***Phosphoprotein enrichment.*** After cell culture treatments and following cell lysis, proteins were enriched in a phosphoprotein enrichment column (Pierce™ Phosphoprotein Enrichment Kit, Thermo Fisher Scientific), according to the manufacturer’s instructions. Briefly, whole cell lysate was applied to a phosphoprotein column containing a proprietary enrichment gel. The samples were incubated in the column for 30 min at 4 °C and washed. Retained proteins were eluted with five column washes with elution buffer. Phosphoprotein content, typically yielding 15–25% of the total protein loaded, were determined using the BCA assay.

*Immunoblotting of mammalian cell extracts.* Total protein was resolved by sodium dodecyl sulphate-polyacrylamide gel electrophoresis (SDS-PAGE) and transblotted using standard procedures. Nitrocellulose membranes (BioRad) were incubated with primary (Supplementary Table 3) and secondary antibodies (Kindle Biosciences, Cell Signaling Technology), and revealed by chemiluminescence using the ECL kit (Millipore Classico or Kindle Biosciences ECL kit) and imaged using a Kwik Quant Imager (Kindle Biosciences).

*Cytotoxicity assays.* Activated caspase-3 and 7 was measured using the Caspase-3/7 Glo kit (Promega). To assess cell viability the Cell Titer Glo® (Promega) assay was used following manufacturer's instructions.

*Drosophila stocks*. Drosophila miR-219 knockout line (miR-219$^{-/-}$) was generated by targeted homologous recombination[8]. MiR-219$^{-/-}$ line was isogenized and backcrossed for more that 10 generations with control line $w^{1118}$ (Exelisis, Inc). Tau-GFP protein trap line was obtained from Bloomington Stock Center (BL-60199. Recombinant line miR-219$^{-/-}$::Tau-GFP was generated in the laboratory for this work. All Drosophila stocks were maintained at 25°C (unless otherwise specified) on a 12-h/12-h light/dark cycle at constant humidity in standard medium.

*Lifespan Assay***.** Lifespan assays were performed as described previously[9]. Briefly, 200 flies per line were sex-segregated within 4 h of eclosion and maintained in small laboratory vials (n = 20 per vial) containing fresh food in a low-temperature incubator at 25 °C and 40% humidity on a 12/12 h dark/light cycle. The flies were then transferred to fresh food vials every 2-3 days and mortality recorded.

*Drosophila brain histology***.** 7, 20 and 45-days old fly heads were fixed in Carnoy's fixative and embedded in paraffin. 6 μm serial frontal sections were prepared through the entire fly brain. Slides were processed and stained with hematoxylin and eosin, and examined by bright-field microscopy. Analysis and quantification of the presence and number of vacuoles was done using the Aperio ImageScope software (Leica Biosystems).

*Immunostaining.* Larval and adult NMJs were dissected under the microscope. Briefly, flies were anesthetized with $CO_2$ and immobilized with dissection pins in Silgar polymere plates with their dorsal side up. Flies were then submerged in PBS containing 4% formaldehyde, allowing the fixing process to start. A dorsal-longitudinal cut was done along the abdomen and dissection pins were used to open the abdomen in a wide-open-book manner. Fat and non-required tissues were removed. After a total fixation process of 20 minutes, abdomens were washed with PBS containing 0,1% Triton-X (PBT) and transferred to 4-well plates for immunostaining. Third instar larvae were dissected similarly and fixed in 4% formaldehyde in PBS, abdomens were washed and incubated with PBT containing 5 % normal goat serum.

Synapses were revealed in larval and adult NMJs with primary mouse monoclonal antibody nc82 against presynaptic BRP protein (DSHB) and anti-HRP rabbit antibody (Jackson ImmunoResearch Laboratories). Samples were then mounted in Vectashield medium with DAPI (Vector Laboratories) after incubation with secondary antibodies anti-mouse Alexa 488 and anti-rabbit Alexa 568 (Invitrogen). Total nc82 positive puncta were quantified in the larval third segment or adult VLM-NMJs (ventral longitudinal muscle neuro-muscular junctions) with Imaris (Bitplane) software.

Adult brains of 7, 20 and 45 days were dissected and fixed in 4% formaldehyde in PBS for 20 min, and then washed in PBS. GFP-tagged drosophila tau was directly visualized.

For apoptotic cell death visualization, primary antibody anti-caspase-3 (Cell Signaling) was incubated overnight in PBT containing 5% normal goat serum. Brains were then washed and incubated with Alexa-568 secondary antibody (Invitrogen), washed again and mounted with Vectashield mounting media containing DAPI (Vector Laboratories). Number of caspase-3 positive cells was counted in at least 10 brains per group with Imaris (Bitplane) software.

Cell death was also visualized in adult brains of male flies of 7, 25 and 45 days old, using terminal deoxynucleotidyl transferase biotin dUTP nick end labeling (TUNEL) labeling according to manufacturer's instructions (DeadEnd™ Fluorometric TUNEL System, G3250 Promega). At least 10 brains were analyzed, and the number of TUNEL-positive cells throughout the entire brain was quantified with Imaris (Bitplane) software.

*Drosophila immunoblotting*. Adult fly heads were homogenized in Neuronal Protein Extraction Reagent supplemented with Halt Protease & Phosphatase inhibitor cocktail (Thermo Scientific). Lysates were incubated on ice for 10 min and centrifuged at 13,000 x *g* for 15 min at 4 °C. Protein concentration was determined using the BCA protein assay (Thermo Fisher Scientific). Samples were resolved by SDS-PAGE and analyzed by immunoblot as previously described[10] with antisera against tau (Dako). Quantification of relative expression was done from three independent experiments using $\alpha$-tubulin (Sigma-Aldrich) as loading control for normalization. The signal intensity was quantified using ImageJ (NIH) software.

*Drosophila QPCR.* Total RNA was isolated from the whole *Drosophila*. Briefly, flies were lysed in QIAzol and homogenized using a QIAshredder column. Total RNA was extracted using the miRNeasy Kit (Qiagen). RNA concentrations were measured with a Nanodrop ND-1000 Spectrophotometer. Equal amounts of RNA were reverse transcribed using the First Strand cDNA Synthesis Kit (Origene) according to the manufacturer's instructions. QPCR for *Drosophila* tau and the endogenous control RpL32 was performed using TaqMan Gene Expression assays (Life Technologies). Real-time PCR reactions were performed in triplicate with MicroAmp optical 96-well plates using a Mastercycler ep realplex (Eppendorf) with the following conditions; an initial step of 10 min at 95 °C, followed by 40 cycles of 15 s at 95 °C, 1 min at 60 °C. Each data point is the result of at least three biological replicates each composed of three technical replicates.

*Sarkosyl extraction.* Sarkosyl extraction was performed, with some modifications, as described[11]. Fifty heads from 20-day-old control ($w^{1118}$) and miR-219$^{-/-}$ flies were homogenized with a motorized mini-pestle grinder in 50μL of homogenization buffer (15mM NaCl, 25mM Tris-HCl at pH 7.4, 1mM EGTA and 1mM EDTA) containing protease inhibitors cocktail (Roche Diagnostics). Homogenized samples were centrifuged at 15000 rpm for 20 minutes at 4°C and the supernatant was stored as soluble fraction. Pellet was resuspended in 50 μL of salt-sucrose buffer (10% sucrose, 0.8M NaCl, 10mM Tris-HCl at pH 7.4 and 1mM EGTA) also containing protease inhibitors, and centrifuged 1 hour at 100.000 x g at 4°C. Supernatant was then incubated with 1% fresh Sarkosyl for 1 hour at 37 °C (5 μL was added per sample), to later be centrifuged for 2 hours at 100.000 x g at 4°C. Pellet was stored as insoluble fraction.

*Associative Olfactory Learning Assay***.** Aversive associative memory was performed as previously described with minor modifications[12]. Male flies 1-3 days old were collected in fresh food vials and stored at 25°C until they reached 7 days old. Flies were then trained and tested in groups of ~ 60 flies. Training and testing were performed under dim red light at 25°C and 70% relative humidity. Flies were exposed to 60 sec of an odor paired with 12 pulses (1.25 sec pulses with 3.75 sec inter-pulse intervals) of electric shocks at 60-V (CS+), followed by 60 sec of a second odor without the shock (CS−), with 45 sec recovery period between odors. 3-octanol (OCT $10^{-4}$ (v/v)) and 4-methylcyclohexanol (MCH $10^{-4}$ (v/v)), diluted in mineral oil, were used as odors. [$10^{-4}$] is a concentration previously tested to show no olfactory attraction or repulsion perception in either fly line. Flies were removed from the setup after the training phase was over, and kept in 25 °C and 70% humidity for 1 hour, when the flies were tested for associative memory. Flies were inserted in the T-maze where they were allowed to choose their preference for one of the odors for a period of 120 sec. Flies were then recollected and counted in order to calculate the Performance Index (PI). PI was calculated as follows: (number of flies that chose the CS− minus the number that chose the CS+)/(number of flies that chose the CS− plus the number of flies that chose the CS+).

### Supplementary Figures

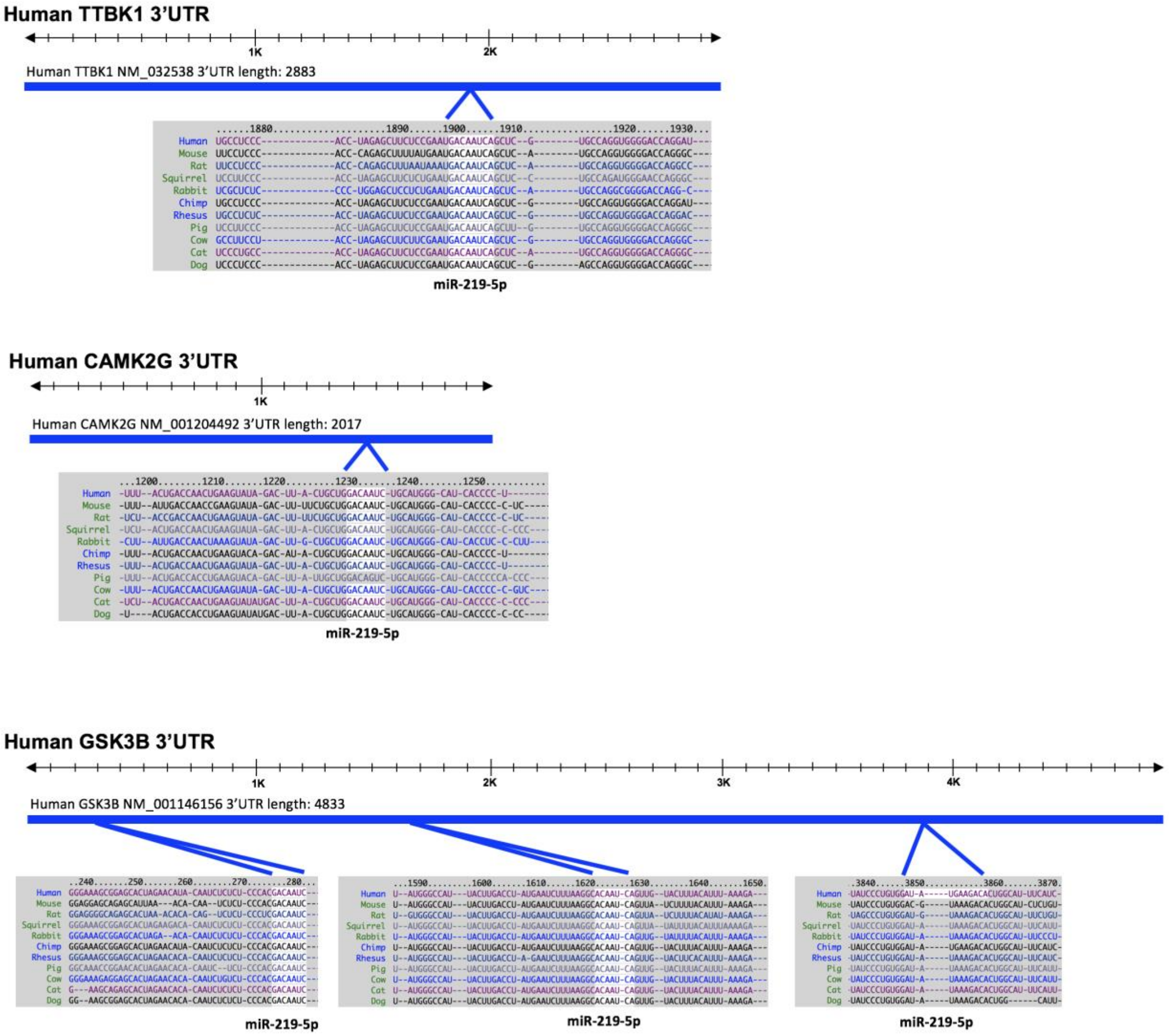


**Supplementary Figure 1**

**3'UTR *in silico* analysis of TTBK1, CAMK2γ and GSK3β.** Predicted miR-219 recognition elements in the 3′-UTR of TTBK1, CAMK2γ and GSK3β using algorithms based on sequence homology[2] are shown.

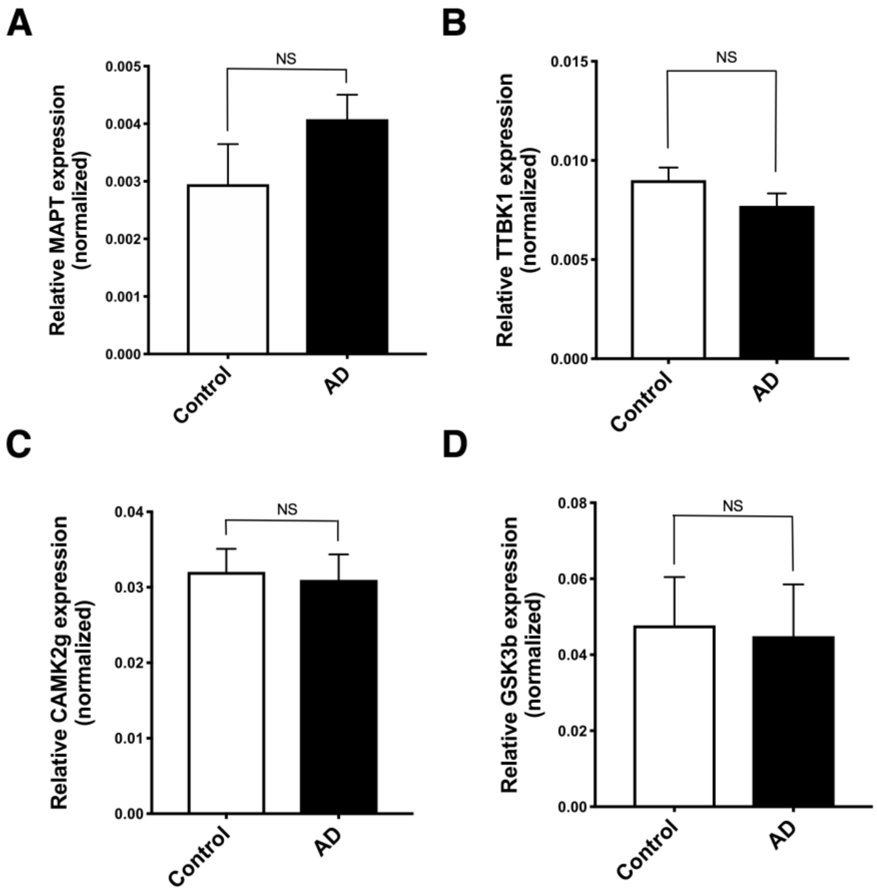


**Supplementary Figure 2**

**mRNA levels of miR-219 targets in Alzheimer disease brains.** qPCR shows that mRNA levels of miR-219 targets Tau (MAPT) (A), TTBK1 (B), CAMK2γ and (C) GSK3β are unchanged in the human postmortem AD (n=26) brain tissue when compared to healthy controls (n=20). NS, P>0.05 by 2-tailed Student's t test.

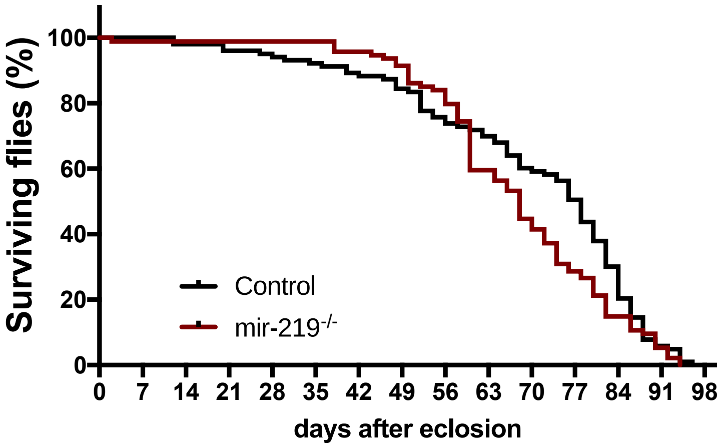


**Supplementary Figure 3**

**Lifespan assays in control (*w*$^{1118}$) and miR-219$^{-/-}$ transgenic flies**. Significant differences in lifespan were not observed between control (*w*$^{1118}$) and miR-219$^{-/-}$. Comparisons of survival curves were performed using the Log-rank (Mantel Cox) test and the Gehan-Breslow-Wilcoxon test.

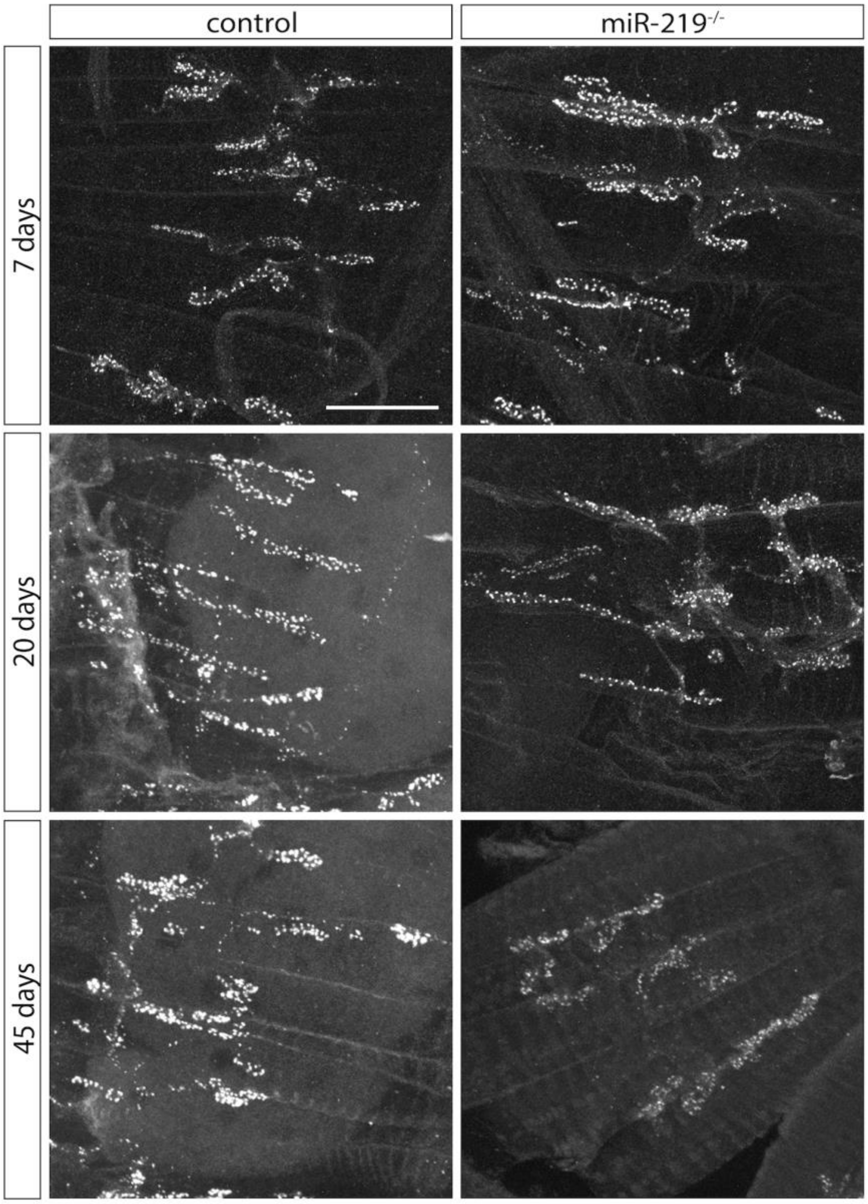


**Supplementary Figure 4**

**NMJs staining of adult Drosophila.** Confocal images of adult NMJs corresponding to the ventral longitudinal muscles in fly abdomens, showing BRP-positive puncta, from indicated genotypes and ages. Scale bar is 20 μm.

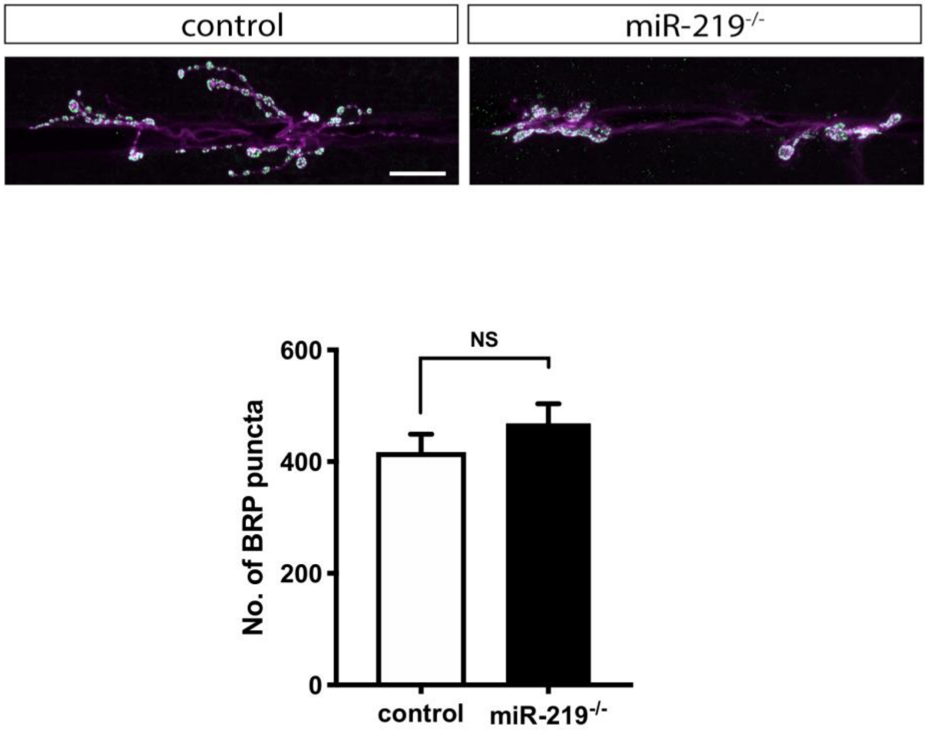


**Supplementary Figure 5**

**Drosophila third instar larval NMJs staining and analysis.** (**A**) Representative images of third instar larval NMJs showing BRP-positive puncta. (**B**) Quantification of the number of synapses showing no significant changes between control and miR-$219^{-/-}$ genotypes (n=7/group). Scale bar is 20 μm. Mean and SEM bars are represented and statistical significance was tested by Mann-Whitney analysis.

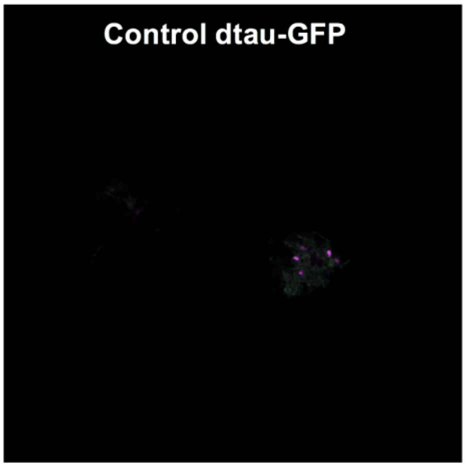


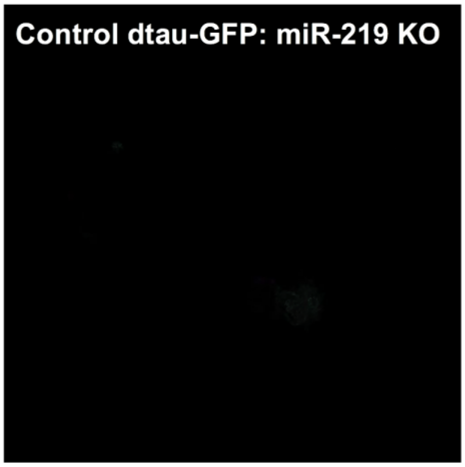


**Video 1** **Video 2**

**Supplementary video 1 and 2**

**Loss of miR-219 results in age-associated tau protein accumulation in the adult brain of Drosophila.** Videos showing confocal stacks of adult brain images from control and miR-219$^{-/-}$ flies expressing GFP-tagged tau. Tau accumulation can be found throughout the brain. Mutant flies lacking miR-219 showed enhanced tau accumulation in different areas of the brain, both in the somas and in the neuropil region (n=3/group).

### Supplementary Tables

**Supplementary Table 1.** Summary of patient brain samples (Entorhinal cortex)

| **Classification** | ***n*** | **Sex (M/F)** | **Average age (year ± SEM)** | **Braak NFT** | **Limbic NFT Frequency** | **CERAD plaque score (0/A/B/C)** | **Clinical diagnosis** |
|---|---|---|---|---|---|---|---|
| Control | 4 | 2/2 | 89 ± 2.9 | 0-2 | Sparse | (3/1/0/0) | Normal |
| AD | 8 | 4/4 | 85.75 ± 12.2 | 4-6 | Moderate-Frequent | (0/0/3/5) | AD |

SEM = standard error of the mean; AD = Alzheimer disease; CERAD plaque score 0 = none, A = sparse, B = moderate, C = frequent

**Supplementary Table 2.** Summary of patient brain samples (BA38)

| Classification | *n* | Sex (M/F) | Average age (year ± SEM) | Braak NFT | Limbic NFT Frequency | CERAD plaque score (0/A/B/C) | Clinical diagnosis |
|---|---|---|---|---|---|---|---|
| Control | 20 | 10/10 | 70.9 ± 1.82 | 0-4 | Sparse-Moderate | (14/4/2/0) | Normal |
| AD | 24 | 6/18 | 80.04 ± 1.21 | 4-6 | Moderate-Frequent | (0/0/4/20) | AD |

SEM = standard error of the mean; AD = Alzheimer disease; CERAD plaque score 0 = none, A = sparse, B = moderate, C = frequent

**Supplementary Table 3**. List of antibodies used in this study

| | Species | Supplier | WB dilution | IS dilution | Epitope specificity |
|---|---|---|---|---|---|
| **Tau antibodies** | | | | | |
| Tau-5 | Mouse | Millipore | 1:1000 | | Total Tau protein |
| Pan-tau | Rabbit | Agilent-Dako | 1:2000 | | Total Tau protein |
| pTau 199/202 | Rabbit | ThermoFisher Scientific | 1:1000 | | Phosphorylation at Ser-199 and -202 |
| pTau 205 | Rabbit | ThermoFisher Scientific | 1:1000 | | Phosphorylation at Thr-205 |
| pTau 231 | Rabbit | ThermoFisher Scientific | 1:1000 | | Phosphorylation at Thr-231 |
| PHF-1 | Mouse | Kind gift[a] | 1:2000 | | Phosphorylation at Ser-396 and -404 |
| **Kinase antibodies** | | | | | |
| TTBK1 | Rabbit | ThermoFisher Scientific | 1:1000 | | Total Tau Tubulin Kinase 1 |
| CAMK2γ | Rabbit | Abcam | 1:1000 | | Total Calcium/Calmodulin Dependent Protein Kinase II Gamma |
| GSK3β | Rabbit | Cell Signaling Technology | 1:1000 | | Total Glycogen Synthase Kinase 3 Beta |
| CDK5 | Mouse | Biocompare | 1:1000 | | Total Cyclin Dependent Kinase 5 |
| **Other antibodies** | | | | | |
| Nc82 | Mouse | Developmental Studies Hybridoma Bank | | 1:20 | Bruchpilot protein |
| Hrp | Rabbit | Jackson ImmunoResearch Laboratories | | 1:200 | Horseradish peroxidase |
| Cc3 | Rabbit | Cell Signaling Technology | | 1:100 | Cleaved Caspase-3, Asp175 |
| β-actin | Mouse | Sigma | 1:2000 | | β-actin |
| α-tubulin (DM1A) | Mouse | Sigma | 1:5000 | | α-tubulin |

[a]Provided by P. Davies, Feinstein Institute for Medical Research, Manhasset, NY 11030; WB, Western blot; IS, Immuno staining
